# Microtubule reorganization during mitotic cell division in the dinoflagellate *Ostreospis* cf. *ovata*

**DOI:** 10.1101/2023.10.17.562701

**Authors:** David Velasquez-Carvajal, Flavie Garampon, Rodolphe Lemée, Sebastian Schaub, Stefania Castagnetti

## Abstract

Dinoflagellates are marine organisms that undergo seasonal explosive proliferation events known as algal blooms. Vegetative cell proliferation is a main contributing factor in these events. However, mechanistical understanding of mitosis and cytokinesis in dinoflagellate remains rudimentary. Using an optimized immunofluorescence protocol, we analysed changes in microtubule organization occurring during the mitotic cycle of the toxic dinoflagellate *Ostreopsis* cf. *ovata.* This study revealed important features of dinoflagellate cell division. We find that the two flagella and the cortical microtubule array persist throughout the mitotic cycle. Two microtubule bundles are present in the cytoplasm originating from the ventral area, where the basal bodies are located: a cortical bundle and a cytoplasmic ventral bundle. The latter associates with the nucleus in the cell centre in preparation for mitosis and with the acentrosomal extranuclear spindle during mitosis. Analysis of tubulin post-translational modifications identifies two populations of spindle microtubules: polar acetylated microtubules whose length is stable throughout mitosis and central tyrosinated microtubules which elongate during chromosome segregation. During cell division a microtubule rich structure forms along the dorsal-ventral axis, associated with the site of cytokinesis, consistent with a cytokinetic mechanism independent of the actomyosin ring typical of animal and yeast cells.

**Summary statement:** Our study describes special features of mitosis and cytokinesis in dinoflagellates and uncovers a new alternative mechanism for cell division, highlighting the plasticity of cell biological process in eukaryotic cells.

## Introduction

Dinoflagellates are unicellular eukaryotes that have adapted to most aquatic ecosystems, from tropical to polar waters. These protists are known to undergo large population expansions that can impact heavily on the marine ecosystem and on human health, a phenomenon known as Harmful Algae Bloom (HAB) (Driscoll et al., 2016). Despite the extensive investigations into the ecological and cellular basis of HABs, the mechanisms controlling dinoflagellate cell proliferation, a major process underlying HAB development, still remain elusive.

Dinoflagellate mitosis, also known as dinomitosis, is unique among eukaryotes. It is a form of closed mitosis, where chromosome segregation occurs within the nucleus, without nuclear envelope breakdown. Differently from yeasts, diatoms and euglenoids, which undergo closed mitosis with a nuclear spindle, in dinoflagellates spindle microtubules are cytoplasmic and traverse the nucleus through tunnels never entering the nucleoplasm (Gavelis et al., 2019). Segregating chromosomes are attached to the nuclear membrane surrounding the nuclear tunnels, through electron dense kinetochore-like structures, which in turn contact microtubules on the cytoplasmic side (Cachon and Cachon, 1977; Oakley and Dodge, 1974). Hence, the interaction between spindle microtubules and segregating chromosomes is indirect, across the nuclear membrane. This configuration poses the question of how the spindle is organized in these organisms and how chromosomes are segregated within the nuclear membrane. EM studies in *Syndinium* (Ris and Kubai, 1974) and in *Amphidinium carterae* (Oakley and Dodge, 1976) showed that the microtubules emanating from the kinetochore-like structures extend exclusively towards the nearby polar region of the spindle and the kinetochore to pole distance remains constant throughout mitosis. Thus, chromosome segregation was suggested to be accomplished by elongation of cytoplasmic microtubules running through the nuclear channels between the two spindle poles

Following nuclear division, the cell divides during cytokinesis. In animal and yeast cells, cytokinesis relies on the constriction of an actomyosin ring. The absence of identifiable myosin-II in many bikonts (Richards and Cavalier-Smith, 2005), including dinoflagellates, however, suggests that an alternative mechanism for cytokinesis must be in place in these organisms. In plants a transient ring of cortical microtubules and actin filaments, the preprophase band, forms early in the cell cycle and marks the site of cytokinesis. During anaphase an equatorial array of microtubules, the phragmoplast, then directs Golgi-derived vesicles to the cortical division site to allow deposition of the new cell wall (Nishihama and Machida, 2001). Mechanisms of cytokinesis that do not involve an actomyosin ring have also been described for many single-celled protists. In *Trypanosoma brucei* cell division occurs by microtubule dependent formation of a unidirectional furrow from the anterior to the posterior of the cell (Sherwin et al., 1989). In *Chlamydomonas reinhardtii* and many related green algae, a microtubule structure, called the phycoplast, forms between the duplicated basal bodies at the anterior of the cell and then grows towards the posterior pole (Ehler and Dutcher, 1998). Actin also accumulates at this site, but treatment with actin depolymerizing drugs does not induce any defect in cell division (Onishi et al., 2020). For dinoflagellates, little is known on how cytokinesis takes place and the contribution of the cytoskeleton to this process. Early EM studies in *A. carterae* (Oakley and Dodge, 1976) and in *Ceratium tripos* (Wetherbee, 1975) showed that cell division occurs by pinching in of the membrane in the equatorial region where microtubules accumulate under the plasma-membrane. Treatment with actin depolymerizing drugs, however, was shown to interfere with completion of cell division in *Prorocentrum micans*, suggesting a contribution of the actin cytoskeleton to cytokinesis, at least in some species (Schnepf et al., 1990).

To gain further insights into the mechanism of dinomitosis and the organization of the mitotic spindle in dinoflagellates, we analysed the spatial and temporal changes in cytoskeletal organization during the mitotic cycle in the dinoflagellate *Ostreopsis* cf*. ovata* (*O.* cf. *ovata*), one of the most common HAB causing species in tropical and subtropical areas (Parsons et al., 2012; Rhodes, 2011) and the most abundant and widely distributed benthic dinoflagellate in the Mediterranean Sea (Ninčević Gladan et al., 2019). *O.* cf*. ovata* cells have a teardrop shape with a pointy ventral side, from which two flagella emanate and a rounded dorsal side, where the nucleus is located during interphase. In interphase cells, a cortical array of microtubules underlies the plasma membrane (cortical microtubule array) and a microtubule bundle runs from the ventral sulcal area towards the inside of the cell (Escalera et al., 2014). Here we show that the cortical microtubule array and the flagella persist throughout the cell cycle. We also analyse changes in the organization of microtubule-based structures during the mitotic cycle and describe novel ultrastructural features of the microtubule cytoskeleton as cells undergo mitosis and cytokinesis. We find that microtubules of the ventral bundle contact the nucleus while it localizes to the centre of the cell in preparation for mitosis. An acentrosomal microtubule-based spindle then forms in the cell centre, transverses the nucleus and separates chromosomes. Analysis of tubulin posttranslational modifications identifies two regions in the anaphase spindle: a stable polar spindle enriched in acetylated tubulin, linking chromosomes to the ventral bundle and a dynamic central spindle, containing tyrosinated tubulin, between separating chromosome masses. In dividing cells, a microtubule-rich structure, free of actin, forms from the dorsal side along the longitudinal axis of the cell where cytokinesis takes place, and divides the cell. Our observations reveal that both mitosis and cytokinesis in *O.* cf*. ovata* are different from yeast and animal cells and support the idea that the mechanisms of cell division are highly diverse outside of opisthokonts.

## Results

### Cortical microtubule array and flagella are maintained throughout the cell cycle

To analyse the changes in the organization of the microtubule cytoskeleton associated with the mitotic cell cycle in *O.* cf*. ovata* we have optimized an immunofluorescence protocol previously developed to stain cortical microtubules (Escalera et al., 2014). We applied it to label microtubules in both cultured cells and wild samples collected in the Bay of Villefranche-sur-mer during summer blooming and obtained similar results with both sources of cells. Staining using the ß-tubulin D66 antibody labelled several structures in the cell (Fig. 1A). Previous description of microtubules in *O.* cf. *ovata* had identified a network of microtubules underlying the plasma membrane in interphase cells (Escalera et al., 2014). We confirmed the presence of this cortical microtubule array (Fig. 1A and movie 1). On the posterior side of the cell, cortical microtubules ran vertically along the dorso-ventral axis covering the entire surface of the cell. On the opposite anterior side, recognized by the presence of the apical pore, instead, cortical microtubules run obliquely from the apical pore to the side of the cell. The anterior and posterior halves are separated by the cingulum, which circles the entire cell (Fig. 1A). The cingulum is delimited by two parallel bands of microtubules, while in the groove of the cingulum tightly packed parallel microtubules run transversally, perpendicular to the lateral bands (Fig. 1A, lateral view). Two previously identified thick bundles of MTs originating from microtubular roots present in the ventral sulcal area, can also be observed: one close to the cell surface on the posterior side of the cell, which we named cortical bundle (CB), and a second deeper in the cytoplasm, herein referred to as ventral bundle (VB, Fig. 1A and movie 1). Specificity of the observed tubulin staining was confirmed by treatment of cells with the microtubule depolymerizing drug colchicine for 1 hour prior to fixation. All microtubules were lost in colchicine treated samples but not in control (dH2O) samples (Fig. 1B).

**Figure 1.**
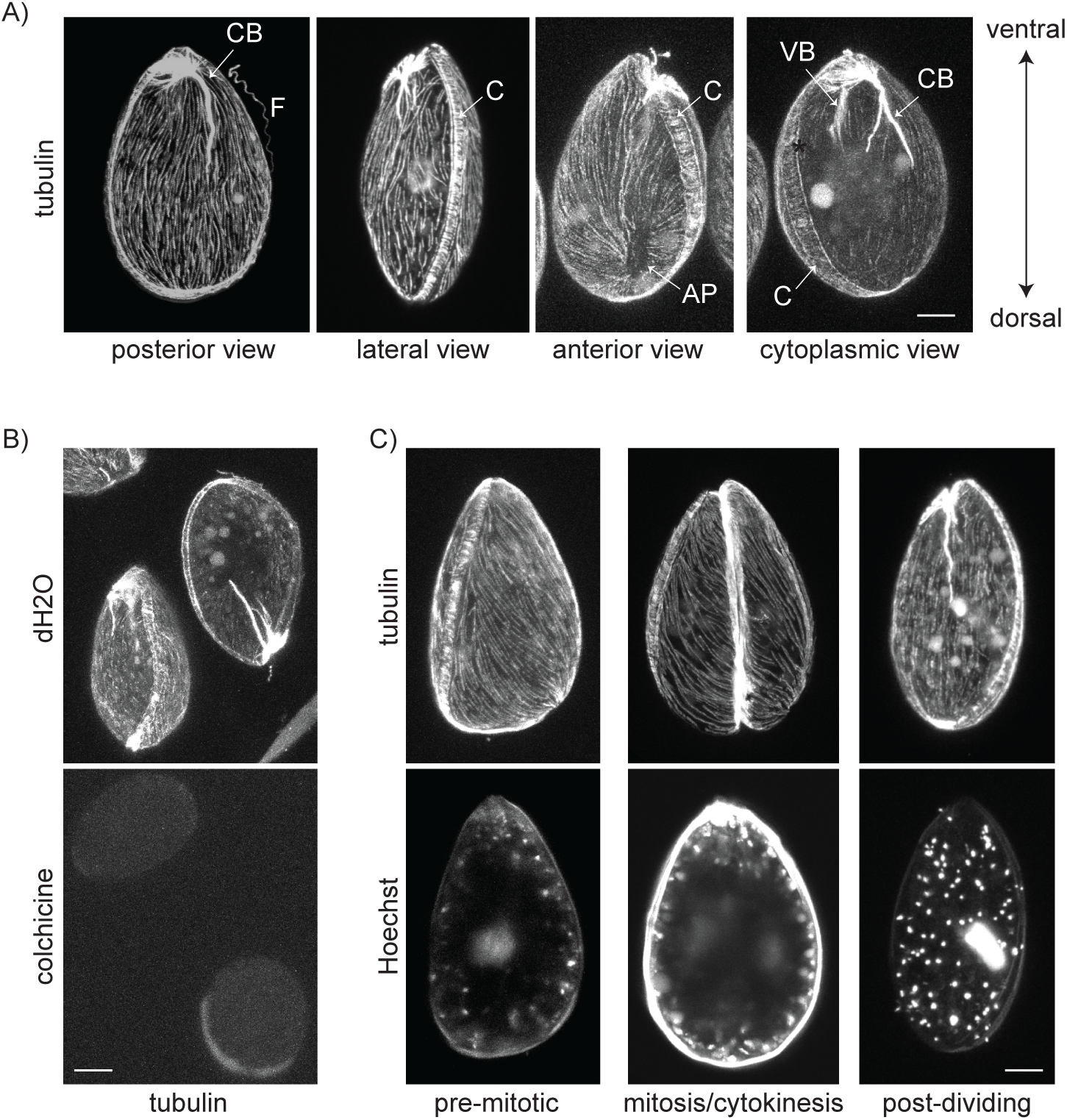
The cortical microtubule array is maintained during mitosis and cytokinesis. **(A)** Representative confocal images of *O.* cf. *ovata* cells stained for ß-tubulin (D66 anti-ß-tubulin antibody). F= flagellum, AP= apical pore; VB= ventral bundle; CB= cortical bundle; C= cingulum. **(B)** Representative confocal images of control (dH2O) and colchicine (2 M) treated *O.* cf. *ovata* cells stained for ß-tubulin (D66 antibody). **(C)** Representative confocal images of pre-mitotic, mitotic and post-dividing *O.* cf. *ovata* cells stained for ß-tubulin (top, D66 antibody) and with Hoechst (DNA, bottom). Hoechst also stains DNA in mitochondria and chloroplasts. All cells are oriented with the ventral side on top, with the only exception of dH2O treated cells in B. All images are projections of the confocal Z-stack covering the whole cell. Scale bars are 10 m.

Using bloom samples collected between 12pm and 4am, which as previously showed are enriched in mitotic and dividing cells (Pavaux et al., 2021), we then asked whether cortical microtubules were maintained throughout the cell cycle. In *O.* cf. *ovata* different cell cycle stages can be identified based on morphological features and on the characteristic position of the nucleus within the cell (Bravo et al., 2012; Pavaux et al., 2021). The nucleus which is located in the dorsal area during interphase (non-dividing cells) relocates to the cell centre in preparation for mitosis (pre-mitotic cells), where the nucleus divides (mitotic cells). Following mitosis binucleated cells divide (cytokinesis) by fission and, using brightfield microscopy a furrow can be observed bisecting the cell to give rise to two post-dividing cells recognized by the presence of the nucleus in lateral position.

We observed that the cortical array of microtubules is stable and persists throughout the cell cycle: in interphase, pre-mitotic, mitotic and post-dividing cells (Fig. 1C). Similarly, flagella, emanating from the ventral pole, were present at all stages of the cell cycle, including mitosis and cytokinesis (Fig. S1). This is in contrast with what was previously described in another dinoflagellate species *Crypthecodinium cohnii* (Perret et al., 1993) or in other algal species like *C. reinhardtii*, where flagella are absent during mitotic cell division (Cross and Umen, 2015; Rosenbaum et al., 1969).

### Changes in microtubule organization during mitosis and cytokinesis

In most interphase cells (> 95%), the two microtubule bundles (VB and CB) extend for a third to half of the cell length towards the cell centre (Fig. 2A, interphase). The cortical bundle is a stable thick bundle of microtubules most likely associated with the flagellar apparatus to anchor the basal bodies to the cell cortex, comparable to the rootlet microtubule of *C. reinhardtii* (Rosenbaum et al., 1969). The ventral bundle instead is a more dynamic structure which changes during the cell cycle. During interphase, microtubules of the VB are compacted for most of the length of the bundle but thin ramifications can be observed extending in the cell centre (Fig. 2A, interphase). In few interphase cells (< 5%), the ventral bundle extends past the cell mid-line and reaches the nucleus in dorsal position (Fig. 2A, interphase). In pre-mitotic cells, when the nucleus is positioned in the cell centre, microtubules branching from the ventral bundle contact the nucleus, forming a structure that resembles an inverted “Y” (Fig. 2A and Fig. S2, pre-mitotic) and enveloping the ventral side of the nucleus.

**Figure 2.**
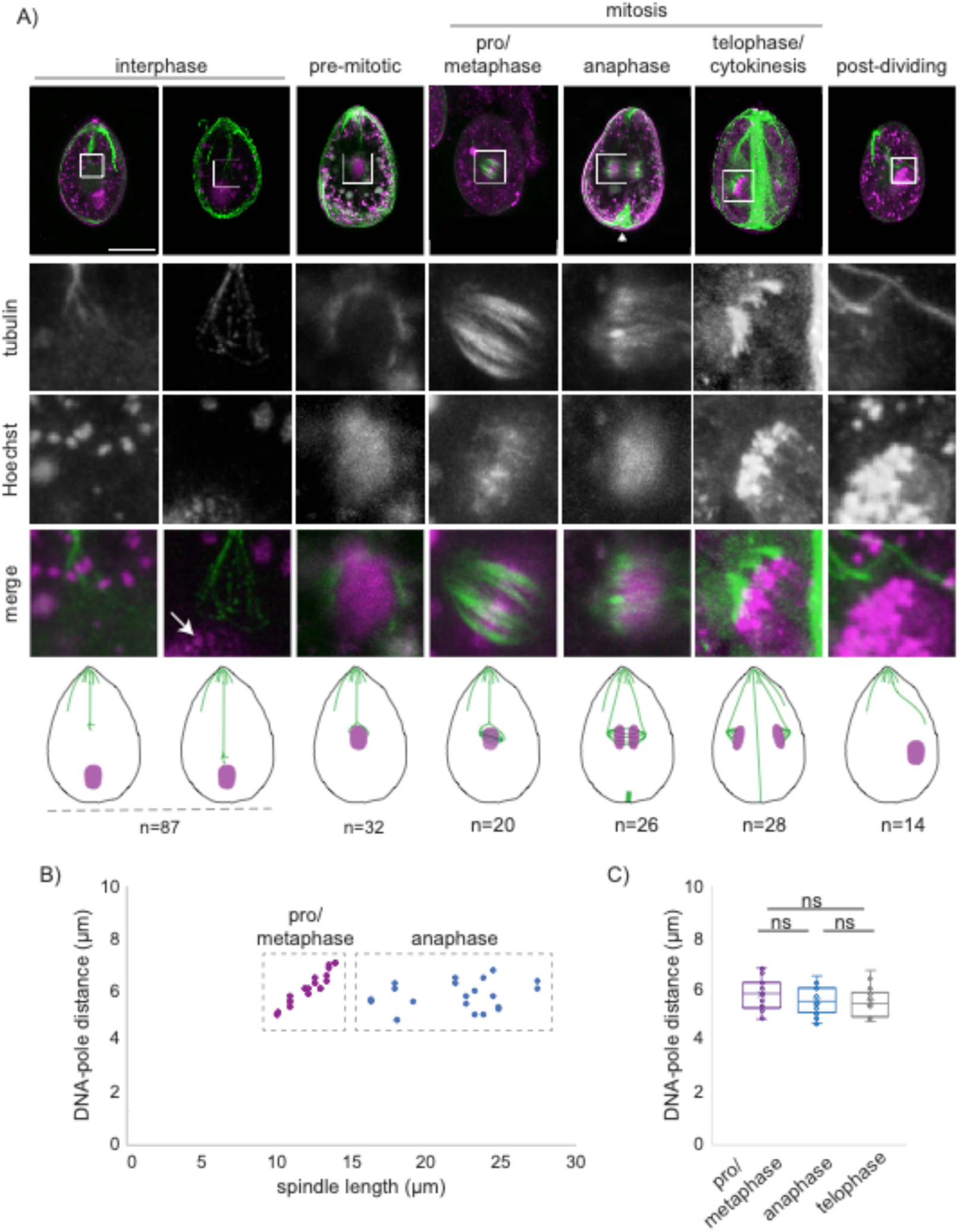
Changes in microtubule organization during dinomitosis. **(A)** Representative confocal images of *O.* cf. *ovata* cells at different cell cycle stages stained with Hoechst (DNA, magenta) and for tubulin (green) using the D66 anti-ß-tubulin antibody. White squares indicate the cell region enlarged underneath each image. All cells are oriented with the ventral side on top. All images are projections of the confocal Z-stack covering the nucleus. Schematic representations of microtubules and nucleus at each analysed cell cycle stage are reported at the bottom. Arrow points to DNA; arrowhead to forming cytokinetic structure. Number of analysed cells (n) is given for each stage. (**B)** DNA-to-spindle pole distance as a function of total spindle length (pole to pole) in pro/metaphase (red) and anaphase (blue) cells. Each dot corresponds to half a spindle. (**C)** Quantification of DNA-to-spindle pole distance in cells at different mitotic stages. Each dot represents half a spindle. Boxes represent 25-75^th^ percentiles, and the mean is shown. T-test was performed using all data points and retrieved non-significant difference (ns). Scale bars are 20 µm.

In mitotic cells, microtubules then organize in 6 to 8 parallel bundles (n= 5 cells) that align along the short axis of the cell to form the mitotic spindle (Fig. 2A and Fig. S2, pro/metaphase), suggesting the presence of 6-8 nuclear tunnels. Spindle microtubules converge at two opposite poles which are connected to the ventral area of the cell by thin microtubule bundles (Fig. 2A), probably originating from splitting of the ventral bundle present in interphase cells. No astral microtubules are visible in mitotic cells. As previously reported for other dinoflagellate species (Bhaud et al., 2000), we observed that in early mitosis, the DNA concentrates in the central region of the spindle onto a metaphase-like plate (Fig. 2A, pro/metaphase), before separating into two discrete masses during anaphase (Fig. 2A and Fig. S2, anaphase). Measurement of pole to pole distance showed that spindle length is constant prior to chromosome segregation (pro/metaphase; average ± SD: 12,1 ± 1,3 µm) and spindle elongation is observed only following chromosome segregation in anaphase when spindle length extends up to 27,5 µm (Fig. 2B). However, we observed that pole to DNA distance, measured as distance between the middle of each DNA mass and the corresponding spindle pole, remains constant during mitosis from prophase to telophase (Fig. 2C; average ± SD: pro/metaphase: 5,95 ± 0,6 µm; anaphase: 5,7± 0,6 µm; telophase: 6,1± 0,5 µm). Thus, chromosome segregation appears to be driven exclusively by extension of the central spindle region.

As shown in Fig. 2A (anaphase, arrowhead), in many anaphase cells, microtubules also accumulate in the dorsal area of the cell. By analysing cells at different mitotic stages, we noticed that a dorsal enrichment in microtubules could be observed already at earlier mitotic stages. This microtubule-rich structure corresponds to the site of cytokinesis (Fig. 2A and Fig. S2, telophase/cytokinesis and Fig. 3A). Three-D reconstruction of cells at different stages of mitosis showed that this microtubule structure develops progressively. In early mitotic cells, up to anaphase, microtubules first arrange in a ring, perpendicular to the cingulum, under the cell surface (Fig. 3A, metaphase/anaphase). As cells progress through anaphase microtubules fill the space between the dividing cells (Fig. 3A, telophase) and by the end of mitosis, when chromosomes are fully segregated to opposite sides of the cell and the central spindle is no longer visible, microtubules fill the entire surface between the two daughter cells, forming a dense microtubule plate (Fig. 3A, cytokinesis).

**Figure 3.**
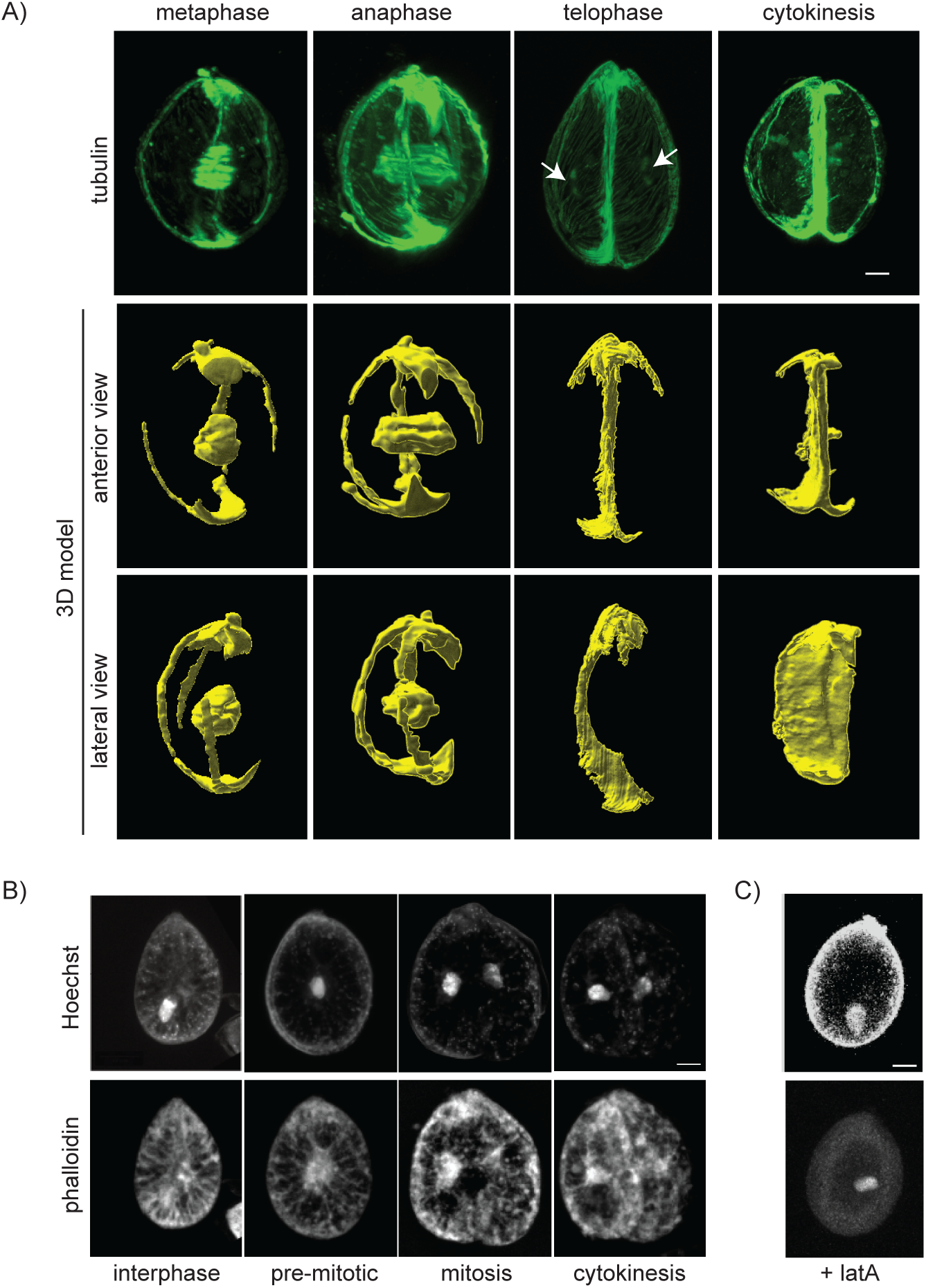
A microtubule-rich plate divides daughter cells in cytokinesis. (A) Confocal images (top) and corresponding 3-D reconstructions (anterior view, middle; lateral view: rotation of anterior view by 90 degrees to the right, bottom) of mitotic cells stained for tubulin using the D66 anti-ß-tubulin antibody. Arrows point to telophase polar spindle. (B) Representative confocal images of *O.* cf. *ovata* cells at different cell cycle stages or (C) of cells treated with the actin depolymerizing drug latranculin A, stained with Hoechst (DNA, top) and with phalloidin (actin, bottom). All cells are oriented with the ventral side on top. Scale bars are 10 µm.

The presence of a microtubule rich plate at the site of cell division suggests that cytokinesis is driven by microtubules in *O.* cf*. ovata*. This is consistent with observations in other protists where microtubule-rich structures drive cell division and are essential for cytokinesis (Ehler and Dutcher, 1998; Francia et al., 2015; Hammarton, 2019; Onishi et al., 2020; Sherwin et al., 1989). Although microtubule enrichment at the cleavage furrow was already observed in dinoflagellates, actin is thought to be required for completion of cell division in some dinoflagellates species, like *P. micans* (Schnepf et al., 1990). We therefore asked whether actin microfilaments are enriched at the cytokinetic plate. Similar to what was previously observed in other dinoflagellate species (Schnepf, 1988; Villanueva et al., 2014), staining with phalloidin showed a diffuse cytoplasmic grid-like pattern (Fig. 3B) which was lost following 5 minutes of treatment with the actin depolymerizing drug latrunculin-A (10 µM, Fig. 3C). Interestingly latranculin A treated cells quickly lost their shape and often become round within 5 minutes of treatment. In untreated cells, we observed actin fibres connecting the cell cortex to the cell centre, where the phalloidin staining is more concentrated. In all cells, irrespectively of cell cycle stage, actin also accumulated around the nucleus (Fig. 3B), as previously described in *C. cohnii* (Perret et al., 1993) and *L. polyedra* (Schnepf, 1988; Stires and Latz, 2018). No discrete actin-containing structure was observed at the site of division, further supporting the existence of a microtubule-based mechanism both for chromosome segregation and for cytokinesis in *O.* cf. *ovata*.

### Different microtubular sub-populations drive cell division

All microtubules are formed by α/ß-tubulin heterodimers whose properties are modulated by post-translational modifications (PTMs) (Bulinski et al., 1988; Janke, 2013; Wolff et al., 1992). Among all identified PTMs, acetylation, tyrosination and poly-glutamylation (Fig. 4A) appear to be conserved across most eukaryotes. We reasoned that analysing tubulin PTMs would allow to distinguish between different sub-populations of microtubules and their changes during the cell cycle, providing insight into how the cytoskeletal structures described above are generated and are related to one another. We therefore analysed tubulin tyrosination, acetylation and poly-glutamylation during *O.* cf*. ovata* cell division (dinomitosis and cytokinesis). We first tested whether these tubulin PTMs could be detected in *O.* cf. *ovata* protein extracts by western blot using specific antibodies. We used the YL1/2 monoclonal antibody which recognizes the C-terminal EEF sequence of yeast α-tubulin (Kilmartin et al., 1982), conserved in *O.* cf*. ovata* α-tubulin; a monoclonal antibody (T6793) which specifically recognizes acetylated lys40 of α-tubulin, and the antibody GT335 against mouse poly-glutamylated tubulin (Wolff et al., 1992). As shown in Fig. 4B, YL1/2 and T6793 antibodies recognized a single protein of the same size as α-tubulin.

**Figure 4.**
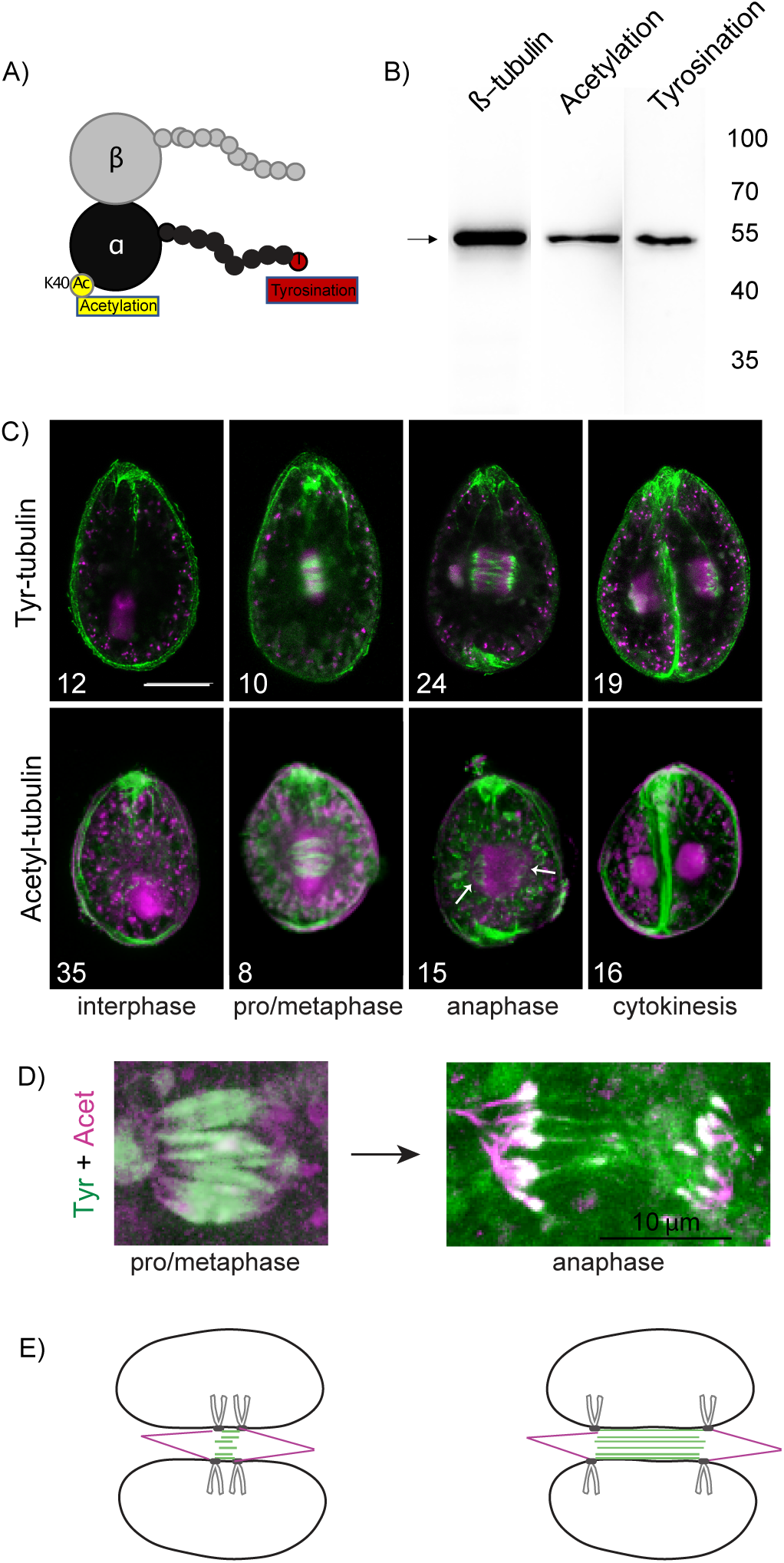
Tubulin post-translational modifications identify different microtubule populations. **(A)** Schematic representation of a α-(black) and β–(grey) tubulin dimer with some posttranslational modifications. **(B)** Western blot analysis of ß-tubulin and tubulin post-translational modifications in whole protein extract from *O.* cf. *ovata* cells. **(C)** Representative confocal images of *O.* cf. *ovata* cells stained with Hoechst (DNA, magenta), and for tubulin with various PTMs (green): tyrosinated tubulin (YL1/2 antibody, top), or acetylated tubulin (T6793 antibody, bottom). The number of analysed cells is indicated on each image. **(D)** Confocal image of *O.* cf. *ovata* pro/metaphase (left) and anaphase (right) spindles co-stained for tyrosinated tubulin (YL1/2 antibody, green), or acetylated tubulin (T6793 antibody, magenta). **(E)** Schematic representation of mitotic nuclei with metaphase and anaphase spindles: chromosomes are in light grey, kinetochore like structures in dark grey, nuclear membrane in black, tyrosinated tubulin in green and acetylated tubulin in magenta. Only 4 chromosomes are represented for simplicity. Scale bars are 20 µm, unless otherwise stated.

Tyrosinated α-tubulin corresponds usually to newly polymerized and dynamic microtubules, whereas removal of the carboxy-terminal tyrosine residue (detyrosination) marks microtubule aging and stabilization (Bulinski et al., 1988; Schulze et al., 1987). Immunolabelling of cycling *O.* cf. *ovata* cells using the YL ½ antibody showed that tyrosinated tubulin is present in the flagella, in the cortical array and in the ventral bundle at all cell cycle stages (Fig. 4C, interphase and Fig. S3A). In mitotic cells both in early (pro/metaphase), and late (anaphase) stages, the YL ½ antibody labelled the central region of the spindle near the midplane, but no signal was detected in the polar spindle regions until telophase (Fig. 4C, mitosis).

Acetylation of lysine 40 (lys40) on α-tubulin instead was shown to be often associated with stable long-lived microtubules (Eshun-Wilson et al., 2019). Immunostaining with the T6793 antibody showed that, at all cell cycle stages, both the cortical array and flagella contain acetylated microtubules (Fig. S3B). Differently from tyrosinated tubulin, in early mitotic cells, before chromosome segregation (pro/metaphase), tubulin acetylation extends over the entire length of the spindle, including the polar region. However, in anaphase cells, following chromosome segregation, acetylated α-tubulin is restricted to the polar region of the spindle which is associated with the DNA (Fig. 4C), but is absent in the central spindle (between the segregating chromosomes). Acetylated-tubulin was also detected at the site of cytokinesis both in the early-cytokinetic ring and in the late-cytokinetic plate (Fig. 4C). Co-staining of *O.* cf. *ovata* cells for acetylated and tyrosinated tubulin did not allow to identify any difference in the cortical array (Fig. S3C), but confirmed that acetylated tubulin is enriched in the polar region of the spindle but absent from the central spindle when the spindle elongates in anaphase (Fig. 4D).

Finally, poly-glutamylation, which is the addition of glutamate side chains of variable length to the carboxy-terminal tail of both α- and ß-tubulin (Bodakuntla et al., 2021; Eddé et al., 1990; RadiBer et al., 1992), is thought to modulate the interaction between microtubules and their partners, MAPs (Microtubule Associated Proteins) and molecular motors, affecting cargo movement, microtubule stability and severing (Bodakuntla et al., 2021). Using the antibody GT335 against glutamylated tubulin (Wolff et al., 1992) we showed that in *O.* cf. *ovata* glutamylated-tubulin is undetectable in non-dividing cells (Fig. S3E, interphase). We could observe tubulin glutamylation exclusively in mitotic cells, starting from early mitosis. As for tyrosinated tubulin, glutamylated tubulin was enriched in the central spindle (Fig. S3E). Differently from acetylated and tyrosinated tubulin, glutamylated-tubulin is not present at the site of cytokinesis.

### Centrin accumulates exclusively at basal bodies

In eukaryotic cells centrioles and basal bodies are essential for the organization of the microtubule cytoskeleton during interphase and mitosis (Wu and Akhmanova, 2017). A well conserved component of both these structures is centrin, a calmodulin-like Ca^2+^-binding protein present in protists, yeast and animal cells. In the green algae *C. reinhardtii*, where it was first identified, centrin localizes to the region of the basal body associated with the flagellum axoneme during interphase, and re-localizes to spindle poles during mitosis to organize the mitotic spindle (Salisbury et al., 1988). As genomic data are not available for *O.* cf. *ovata* we generated transcriptomic data from cultured *O.* cf. *ovata* cells (strain MCCV54) during exponential growth and searched for centrin homologues. Analysis of this transcriptomic dataset allowed the identification of a centrin transcript in *O.* cf. *ovata* with 55% identity to human centrin and 53% to *C. reinhardtii* centrin (Fig. 5A). Using a polyclonal antibody against *C. reinhardtii* centrin (20H5), we then sought to identify microtubule organizing centres (MTOCs) in *O.* cf. *ovata* cells. In western blot assays the 20H5 antibody recognizes a single *O.* cf. *ovata* protein of the expected (20kDa) molecular weight (Fig. 5B). In cells the 20H5 antibody intensely labelled a structure in the ventral region of the cell, near the flagella insertion site where the basal bodies are located (Fig. 5C). In line with our observation that flagella are maintained during the entire cell cycle (Fig. S1), we observed that centrin, which is required for flagellar motility (Fenchel, 2001), was maintained in the ventral area throughout mitosis and cytokinesis. Consistently, we did not observe accumulation of centrin at spindle poles at any stage of mitosis, supporting earlier EM studies suggesting that dinoflagellate spindles are acentriolar. The centrin antibody also stained fibrous structures which extend from the basal bodies towards both the cell centre and the cell cortex (Fig. 5C).

**Figure 5.**
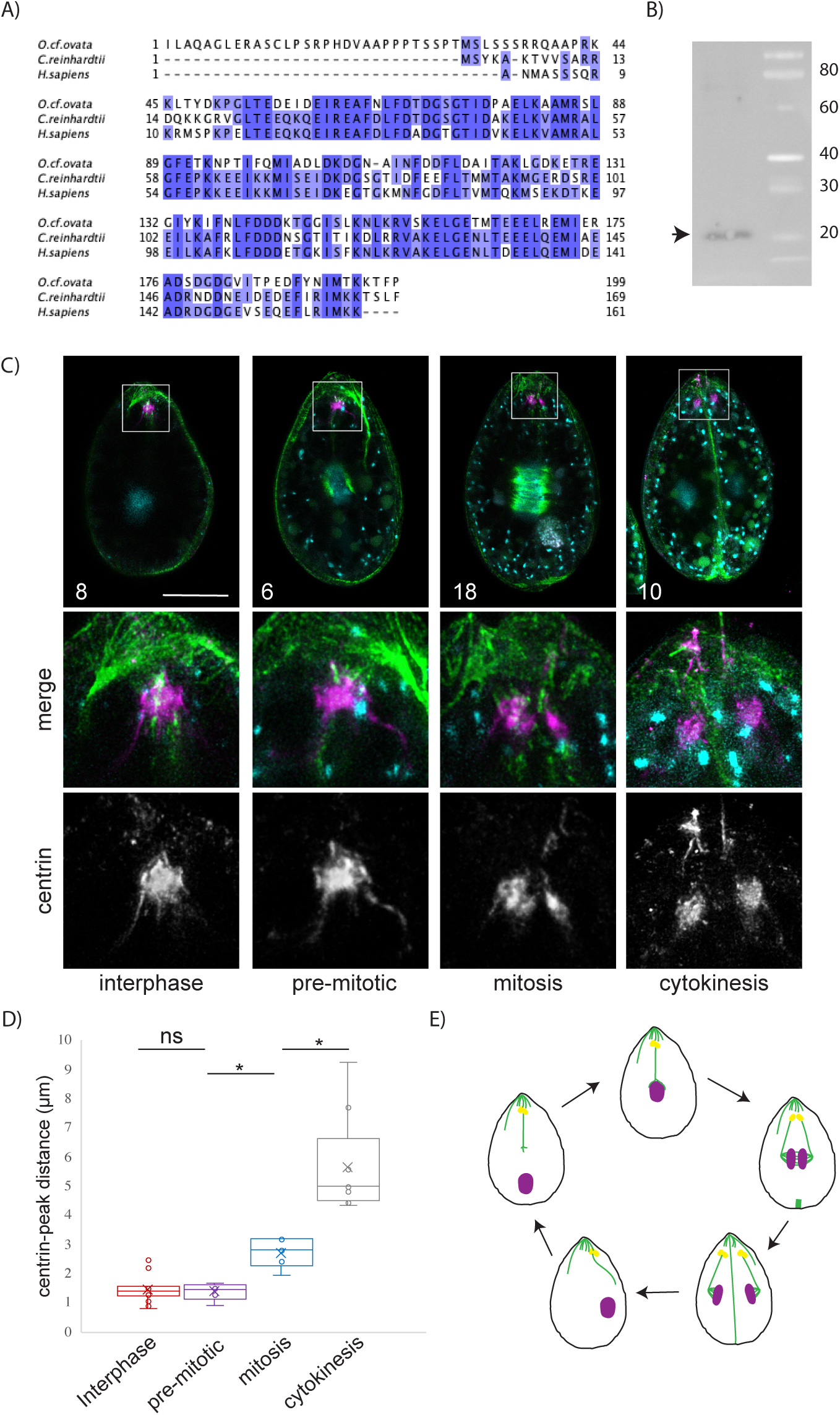
Centrin localizes to basal bodies throughout the mitotic cycle. **(A)** Alignment of centrin sequences from *O.* cf. *ovata, C. reinhardtii*, and *Homo sapiens*. Coloured residues (blue) are identical across species. **(B)** Western blot analysis of centrin in whole protein extract from *O.* cf. *ovata* cells. **(C)** Representative confocal images of *O.* cf. *ovata* cells stained with Hoechst (DNA, cyan), for tubulin using the YL1/2 rat anti-7-tubulin antibody (green) and for centrin using the mouse 20HA antibody against centrin (magenta). White squares indicate the cell region enlarged underneath each image. All cells are oriented with the ventral side on top. All images are projections of the confocal Z-stack covering the nucleus. **(D**) Quantification of distance between peaks of fluorescence corresponding to centrin staining at different cell cycle stages. T-test was performed using all data points and retrieved either non-significant difference (ns) or a p value < 0,001 (*). **(E)** Schematic representation of changes in nuclear and basal body position and microtubule organization during the mitotic cell division cycle of *O.* cf. *ovata*. Scale bars are 20 µm.

Based on our observations we cannot determine when basal body duplication takes place. However, as shown in Fig. 5C we observed that the two centrin-labelled basal bodies are juxtaposed in interphase, separate in mitosis and then segregate to the two daughter cells in cytokinesis. Quantification of the distance between the two peaks of fluorescent intensity in centrin-labelled cells confirmed this observation (Fig. 5D), showing that the two basal bodies separate when the spindle forms as the distance between the two peaks remains constant in interphase (average ± SD: 1,4 ± 0,4µm) and pre-mitotic cells (1,4 ± 0,3µm) but increases to 2,7 µm (± 0,5 µm) in mitotic cells containing a fully formed spindle. In telophase the cytokinetic plate then forms between the two fully segregated basal bodies (5,6 ± 1,7 µm), generating two daughter cells with their own flagellar apparatus.

## Discussion

Due to the unique characteristics of mitosis in dinoflagellates, the term dinomitosis has been used to describe how chromosomes are segregated in those organisms. Here, we have analysed cytoskeletal changes associated with dinomitosis and cytokinesis in the dinoflagellate *O.* cf. *ovata*. which are summarized in Fig. 5 E. In interphase cells the nucleus is located near the dorsal side of the cell and does not interact with microtubules of the ventral bundle. Cortical microtubules run under the plasma membrane and as previously shown for another dinoflagellate, *C. cohnii* (Perret et al., 1993), they persist throughout the cell cycle, including mitosis and cytokinesis. This is similar to what was observed in other unicellular organisms, such as *Trypanosoma brucei*, where the subpellicular microtubule corset which is required for cell shape (Sinclair et al., 2021), is maintained during cell division and is only remodelled locally at the site of furrow formation during cytokinesis (Gull, 1999).

The first sign of preparation for mitosis in *O.* cf. *ovata* is the re-localization of the nucleus to the centre of the cell. We observed that while in the cell centre, the nucleus is always associated with microtubules of the ventral bundle and in few cells this association is also observed while the nucleus is in dorsal position. This configuration suggests that nuclear re-positioning is microtubule based and that extension and branching of the microtubule bundle towards the dorsal side of the cell might be a pre-requisite for nucleus-microtubule interaction and could be considered as an early indicator of preparation for mitosis. Centrin filaments are also associated with the ventral bundle and with the basal bodies where the ventral bundle originates from. It is therefore possible that nuclear movement relies not only on the mechanical action of microtubules and associated motors but also on the contractile activity of centrin, as already observed in nuclear repositioning during mitosis in *C. reinhardtii* (Salisbury et al., 1988). In this organism the two flagella are reabsorbed prior to division and during prophase two major microtubule fibres extend from the basal bodies toward the nucleus and reposition it from the middle of the cell to the apical side, where the basal bodies are located. Centrin which forms filamentous structures that link the basal bodies to the nucleus (Geimer and Melkonian, 2005; Wright et al., 1989) provides contractility to the microtubule bundles and contraction of centrin fibres at mitotic entry allows nuclear movement towards the apical cell surface (Salisbury et al., 1988). This movement is followed by separation of the basal bodies in prophase. Indeed, we observe a similar separation of centrin-labelled bodies in early mitotic stages in *O.* cf. *ovata*.

Upon localization of the nucleus to the cell centre, the mitotic spindle forms, with 6-8 spindle microtubule bundles traversing the nucleus along the short axis of the cell, in line with previous observations in other gonnyalucoid species, the dinoflagellate clade with the highest number of nuclear tunnels (Gavelis et al., 2019). Throughout mitosis, the spindle remains connected to the cell ventral cortex through two microtubule fibres that probably originate from splitting of the ventral bundle. This configuration is similar to the three-pronged fork described in *C. cohnii* where the spindle poles are connected to the basal bodies by a bundle of microtubules, the desmoses (Perret et al., 1993). In animal, yeast and several algal cells, spindle formation is usually driven by the centrosomes localized at spindle poles. Although more markers will need to be analysed, the geometry of the spindle with the association of spindle poles to the microtubules of the ventral bundle and the absence of centrin accumulation at spindle poles suggests that *O.* cf. *ovata* spindles are acentrosomal. Indeed, EM studies in other dinoflagellate species have never identified centrioles in the vicinity of the nuclear membrane, with the only exception of Syndiniales (Bhaud et al., 2000; Drechsler and McAinsh, 2012; Moon et al., 2015). Acentrosomal spindle formation has been observed in animal and plant cells. In those cells spindle organization is driven by chromosomes, often through the formation of a local gradient of RanGTP (Deng et al., 2007; Lee and Liu, 2019). However, this is difficult to envisage for dinomitosis as spindle microtubules are cytoplasmic and indirectly contact the chromosomes which are connected to the inner side of the nuclear membrane.

Our analysis of microtubule PTMs allowed the identification of two regions in anaphase spindles with different characteristics: a polar region highly acetylated and constant in length (constant pole-DNA distance), which is anchored to the ventral area by the ventral bundle, and a central spindle region which is instead enriched in tyrosinated tubulin and whose length increases during mitosis (Fig. 4D,E). Similar to kinetochore microtubules in animal spindles (Wilson and Forer, 1989), the stable acetylated microtubules link chromosomes to spindle poles and most likely correspond to the k-fibers observed by EM both in other dinoflagellates (Oakley and Dodge, 1976; Ris and Kubai, 1974) and in the flagellate hypermastigote *Barbulanympha* (Inoué and Ritter, 1978; Ritter et al., 1978). Different from most animal spindles, instead, the interpolar microtubules running between opposite spindle poles, correspond to the tyrosinated microtubules described here. Interestingly the tyrosinated microtubules never reach the spindle poles. In anaphase spindles, tyrosinated microtubules appear to overlap with acetylated microtubules only in the vicinity of the segregating chromosomes, possibly at the level of the kinetochores. Based on these studies we suggest that during pro/metaphase of dinomitosis mitotic chromosomes become anchored to opposite spindle poles by stable acetylated polar microtubules. During anaphase polymerization of the more dynamic tyrosinated interpolar microtubules then drives chromosome segregation (anaphase B) without prior poleward chromosome movement towards the spindle poles, thus suggesting a lack of anaphase A. The exact organization of the central microtubule array as well as the nature of the molecular motors generating the pushing force remains unclear. Further studies will be required to determine whether chromosome segregation relies on microtubule polymerization or sliding of interdigitating central spindle microtubules. Either way, this mechanism is reminiscent of the central-spindle driven chromosome segregation described during meiosis in *C. elegans* oocytes (Laband et al., 2017) and during mitosis in *C. elegans* embryos without centrosomes (Cowan and Hyman, 2004). In both cases, kinetochore microtubules act as a scaffold to anchor chromosomes to spindle poles and correctly localize them onto the metaphase plate during spindle polarization (Pitayu-Nugroho et al., 2023), while polymerization of microtubules of the central spindle generates an outward pushing force to allow chromosome segregation. The presence of this mechanism in distantly related nematodes and dinoflagellates suggests that it might have emerged early in evolution and could represent a primitive microtubule-based mechanism for chromosome segregation which predates the evolution of cortical interactions to generate pulling forces and ensure higher accuracy in chromosome segregation. Indeed, a similar mechanism can also be observed in bacteria where segregation of low copy number plasmids relies on the polymerization of antiparallel ParM filaments, whose elongation pushes plasmid clusters towards the poles of the cell, allowing their segregation before cell division (Gayathri et al., 2012).

Starting from anaphase, a microtubule-rich structure is formed from the dorsal side, opposite the basal bodies. This structure coincides with the cleavage plane, suggesting a direct implication for microtubules either in placement of the cleavage plane and/or cytokinesis. This observation is in line with the idea that actomyosin driven cytokinesis is a novel acquisition of unikont (animal, yeast and amoeba) cells. Indeed, the lack of Myosin II in most bikont species (Richards and Cavalier-Smith, 2005), including dinoflagellates, supports the use of alternative mechanisms for cytokinesis. In many unicellular protists and in plant cells microtubules have a central role in cytokinesis and form specialized structures associated with the site of cytokinesis. In plants phragmoplast microtubules form in the centre of the cell and then expand centrifugally towards the cell membrane. In *C. reinhardtii* and in other related green algae, instead, cytokinesis is associated with a special array of microtubules, the phycoplast, which forms at the anterior end of the cell where the microtubule rootlets are located and grows unidirectionally towards the opposite pole of the cell (Ehler and Dutcher, 1998). The cytokinetic plate we described here for *Ostreopsis* bears similarities with the algal phycoplast as it forms at the dorsal side and then grows in a unidirectional fashion. However differently from *C. reinhardtii*, the cytokinetic plate described here originates opposite to the basal bodies and grows towards them, suggesting that, as already observed for the plant phragmoplast, the mechanism of microtubule nucleation in the dinoflagellate cytokinetic plate is most likely independent of centrosomes and basal bodies.

## Materials and Methods

### Cell culture and bloom sampling

Strains of O. cf. *ovata* (MCCV054) obtained from the Mediterranean Culture Collection of Villefranche-sur-Mer (MCCV), EMBRC-France, were cultured in autoclaved and filtered enriched seawater, at a salinity of 38 g/L, as described by Guillard and Hargraves (L1 enriched seawater medium) (Guillard and Hargraves, 1993). Cultures were incubated at 22 °C under a 14:10 light/dark cycle (250 µmol m-2 s-1). Cells were fixed for staining during the proliferative phase (day 6 to 10) in order to obtain the greatest number of cells in division.

Bloom samples were collected in Villefranche-sur-mer, on the Mediterranean French coast (43°41′34.83″ N and 7°18′31.66″ E), where yearly blooms occur. Samples used for immunofluorescence were collected in 2018, 2019 and 2020 between 12:00 pm and 4:00 am (Pavaux et al., 2021).

### Actin staining and immunofluorescence

For immunofluorescence a previous published protocol was modified (Escalera et al., 2014). Cells were fixed in cold fixative solution (80% methanol, 10% DMSO (Sigma-Aldrich) and 0,5 mM EGTA pH 7.5 (from 0,5 M EGTA pH 7.5 stock solution in distilled water)) for at least 24 hours, at -20°C to improve membrane permeabilization. Samples could be maintained in fixative solution for several months. When needed, samples were rehydrated progressively in methanol-PBS (60%, 40%, 20%) and finally in PBS, with a 10 minutes incubation at each step. Cells were then permeabilized by two consecutive incubations in PBT (PBS with 0.1% Triton X-100) for 15 minutes at room temperature (RT). Cells were recovered by centrifugation, 5 minutes at 0.2 RCF, and then blocked for 30 minutes at RT in PBT containing 5% BSA (PBT-BSA) and then incubated at 4°C in PBT-BSA containing primary antibody at the appropriate dilution for 3-6 days to maximize antibody penetration. With incubations shorter that 2 days structures deep into the cytoplasm were not labelled. For microtubule staining, D66 anti β-tubulin antibody (against tubulin from *Lytechinus pictus*, mouse, Sigma-Aldrich) was used at 1:400 dilution; for acetylated tubulin, T6793 antibody (Clone 6-11B-1, mouse, Sigma-Aldrich) was used at 1:400 dilution; for tyrosinated tubulin, YL1/2 antibody (ab6160; rat, Abcam) was used at 1:400 dilution; for polyglutamylated tubulin, monoclonal antibody anti polyglutamylated (clone GT335, mouse, Adipogen); the anti-centrin antibody (clone 20H5; mouse, Merk) was used at 1:100 dilution.

After incubation with the primary antibody, cells were washed three times in PBT, and then incubated over-night at 4°C in PBT-BSA with the appropriate secondary antibody at a 1:400 dilution (fluorescently-conjugated anti-mouse and anti-rat antibodies, Jackson ImmunoResearch). After over-night incubation, cells were washed twice with PBS, and then incubated for 10 minutes with 1µg/ml Hoechst-33342 (Sigma-Aldrich) to label the DNA, rinsed twice with PBS and finally resuspended in 100-200 µl of PBS. 50 µl of cells were mixed with 50 µl of citiflour AF1 (Science Services) and mounted on slides for imaging. Each experiment was repeated 3-5 times, 25-50 cells were counted for each sample and 5-15 images were collected for further analysis.

Actin filaments were visualized using CytoPainter Phalloidin-iFluor 488 Reagent (ab176753, Abcam). Cells were fixed in 4% paraformaldehyde in PBS for at least 24 hours at 4°C. Cells were then washed three times in PBS and permeabilized as described for immunofluorescence. After a 30-minute incubation in PBT-BSA, cells were then incubated in PBT containing phalloidin at 1:400 dilution, for 1 hour at RT in the dark. Cells were then washed twice in PBS, and mounted in citiflour AF1 (Science Services) for immediate imaging.

Images were acquired with Leica SP8 and Stellaris 5 microscopes both equipped with a 63X/1.4 Oil and a 40X/1.1W objective, which are part of the Plateforme d’Imagerie Microscopique (PIM) of the Institute de la mer de Villefranche sur mer. On stellaris 5 images were deconvolved with Lightning. When present, Z-Drift was corrected using a custom-made tool (https://github.com/SebastienSchaub/SatAndDriftEstimator). Spindle measurements and image analysis were performed in Fiji (49) with home-made 3D distance tool (https://github.com/SebastienSchaub/3DtDist) and visualization on Imaris (Imaris, BitPlane, Zurich, Switzerland).

### Protein extraction and western blot analysis

To test antibodies, proteins were extracted from cells of strain MCCV-054. 50 ml of cells were grown in L1 media and collected 14 days after inoculum by filtration through Durapore 5 µm PDVF membranes (Millipore) using a manifold system. Cells were recovered from the filters in 1 ml of PBS and then pelleted by centrifugation at 5000rpm for 5 minutes. Pellets were then fast frozen in liquid nitrogen, then resuspended in an equal volume of 2x Laemmli (100 mM Tris-HCl at pH 6.8, 4% SDS, 0.2% bromophenol blue, 20% glycerol, 200 mM dithiothreitol) and heated at 100°C for 5 minutes. Samples were separated on a 10% SDS-polyacrylamide gel and then transferred to a nitrocellulose membrane. Membranes were blocked for 1h in Tris Buffer-Saline (TBS: 20 mM Tris Base, 150 mM NaCl) containing 2% milk, 0.1% tween-20 (TBS-Tween-milk) and then incubated in blocking solution containing the appropriate antibody overnight at 4°C. For centrin detection the 20H5 antibody (Merck) was diluted 1:2000, for detection of tubulin (DM1a antibody, Sigma-Aldrich, mouse) and of PTMs (tyrosinated tubulin: YL ½ antibody, ab6160; rat monoclonal, Abcam; acetylated tubulin: T6793 antibody, clone 6-11B-1, mouse, Sigma-Aldrich; polyglutamylated tubulin: GT335, mouse, Adipogen) all antibodies were diluted 1:3000. Following antibody incubation, membranes were washed 3 times with TBS-Tween (TBS with 0.1% tween-20) and incubated with an appropriate horseradish peroxidase-conjugated secondary antibody (Jackson ImmunoResearch) at 1:10000 dilution in TBS-Tween-milk for 1 hour at RT. After 3 washes with TBS-Tween, signal detection was carried out using the SuperSignal West Pico chemio-luminescent substrate (Thermo Scientific) as described by the manufacturer and imaged in the Vilmen-Loubet Fusion machine.

### RNA extraction, transcriptome annotation and sequence analysis

For RNA sequencing total RNA was extracted from cells of strain MCCV-054. 150 ml of cells were grown in L1 media and collected 6, 12 and 18 days after initial inoculum as for protein preparation.

Filters with cells were placed in 1ml of TRI-reagent (Sigma-Aldrich) and incubated on ice for five minutes. Cells were broken by beating in the presence of ∼ 3 g of glass beads (Sigma) in a 220 V bead beater at 4000 rpm for 40s. RNA was then extracted using a RiboPureTM (Ambion) kit following the manufacturer’s instructions. Residual DNA was removed by treatment with DNase I (Q1 DNAse, Promega) for 1h at 37 °C (2 units per sample) followed by purification with the RNeasy miniElute Cleanup kit (Qiagen).

RNA purity, quantity and integrity were assessed for each sample using a Nanodrop ND-1000 and 2100 Bioanalyser. RNA samples were then sent to BGI-Tech, China for library generation and sequencing with the DNBSEQ™ technology platforms to generate 100 base pair (bp) paired end reads.

*De novo* assembly of reads was carried out with Trinity version 2.4.0 (Grabherr et al., 2011) using default parameters. The full assembly was filtered to the longest transcript per gene group making use of the ’_i’ designations of the Trinity transcript identifiers. Open reading frames were predicted using TransDecoder v5.3. These protein sequences were searched against the NCBI NR database and sequences whose best hit was to Dinophyceae were retained. To annotate these sequences, the pantherScore2.1.pl script was used to search them against the PANTHER hidden Markov model library (version 14.1). Multiple sequence alignments were performed using Clustal Omega and viewed using Jalview Vs 2-a (Waterhouse et al., 2009).

## Acknowledgments

We are grateful to L. Besnardeau for technical support, to E. Christians, S. Marro and M. Hagström for discussion and sharing of reagents, to J. Chenevert and R. Copley for critical reading of the manuscript. We thank the imaging platform, PIM (member of the MICA microscopy platform) where image acquisition was conducted. PIM is supported by EMBRC-France, whose French state funds are managed by the ANR within the investments for the future program under reference ANR-10-INSB-02. DV was supported by Minciencias-Colombia - financial support 783, PhD program.

