## Supplemental Figure 1-3 for "Microtubule reorganization during mitotic cell division in the dinoflagellate *Ostreospis* cf. *ovata*"

### Supporting information

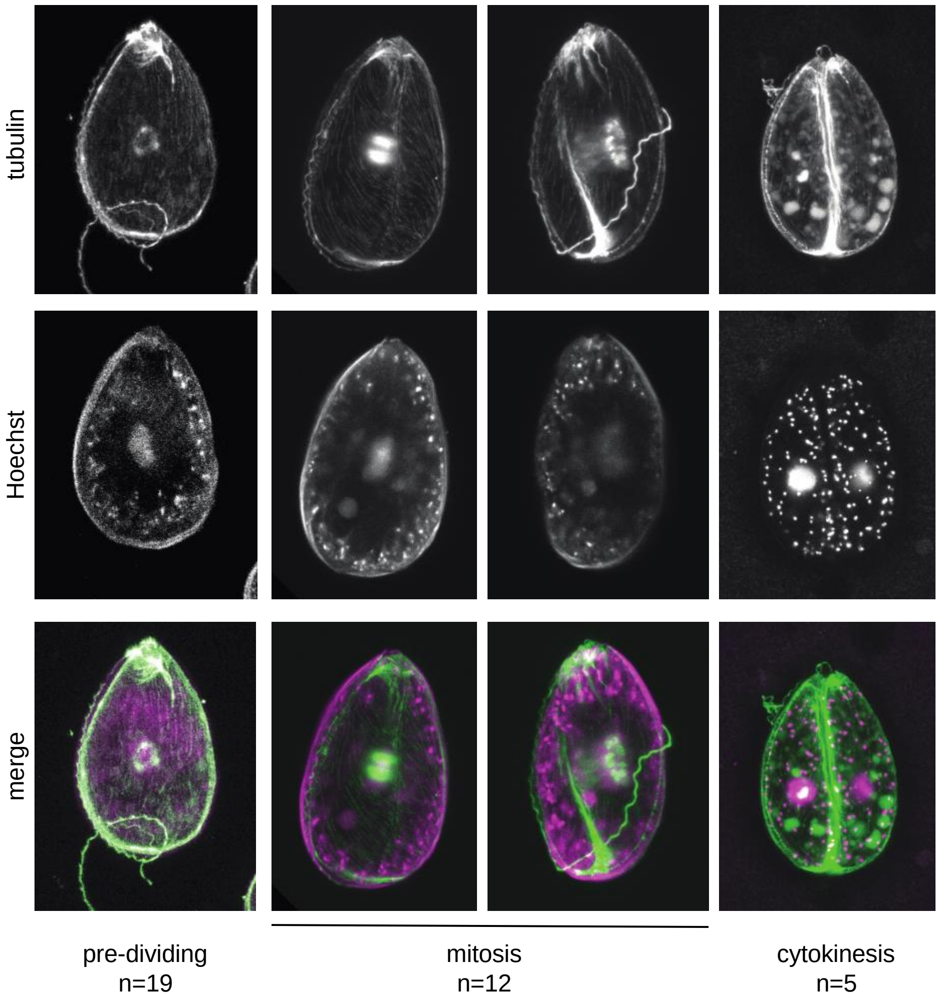

### Figure S1: Flagella are maintained during mitosis and cytokinesis

Representative images of pre-mitotic, mitotic, cytokinetic and post-dividing cells stained for microtubules (anti-tubulin, green) and for DNA (Hoechst, magenta). Images are Z-projections of confocal stacks spanning the whole cell. Scale bar is 10 µm.

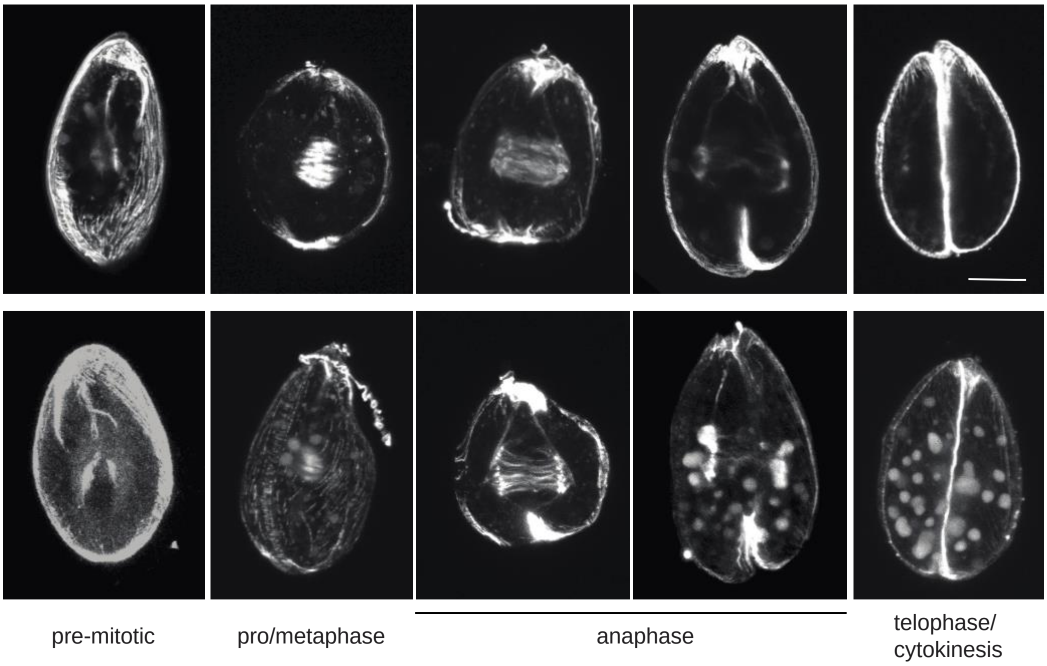


**Figure S2: Microtubule re-organization during mitosis**

Representative images of pre-mitotic, pro/metaphase, anaphase and telophase/cytokinetic cells stained for microtubules (anti-tubulin). Images are Z-projections of confocal stacks spanning the nuclear region. Scale bar is 10 μm.

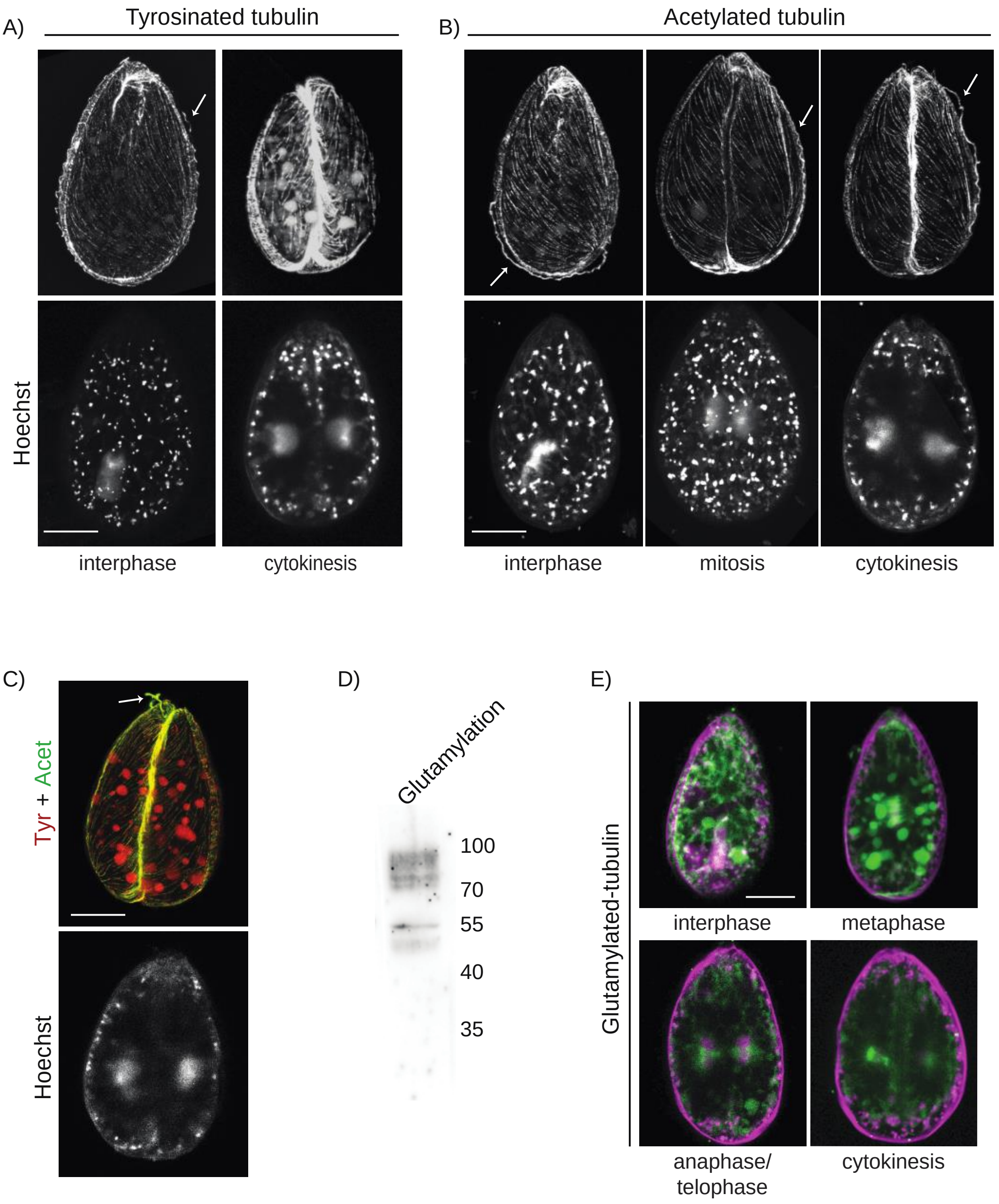
A)
Tyrosinated tubulin
B)
Acetylated tubulin
Hoechst
interphase
cytokinesis
interphase
mitosis
cytokinesis
C)
Tyr + Acet
Hoechst
D)
Glutamylation
100
70
55
40
35
E)
Glutamylated-tubulin
interphase
metaphase
anaphase/
telophase
cytokinesis

**Figure S3: Post-translational modification of flagellar and cortical microtubules during the cell cycle.**

**A)** Representative images of cells at different cell cycle stages (indicated under each image) stained for tyrosinated tubulin and **B)** acetylated tubulin. Arrows point to flagella. **C)** Cytokinetic cell stained for tyrosinated (red) and acetylated (green) tubulin (top) and Hoechst (DNA, bottom). **D)** Western blot analysis of poly-glutamylated tubulin in whole protein extract from *O.* cf. *ovata* cells. **E)** Representative images of cells at different cell cycle stages (indicated under each image) stained for Glutamylated tubulin (green) and Hoechst (DNA, magenta). Scale bars are 20 µm.

**Movie 1:** 3D reconstruction of *O.* cf. *ovata* interphase cell from confocal acquired z-stack, with microtubules in green and DNA in magenta.
